# Glutamate transporters contain a conserved chloride channel with two hydrophobic gates

**DOI:** 10.1101/2020.05.25.115360

**Authors:** Ichia Chen, Shashank Pant, Qianyi Wu, Rosemary Cater, Meghna Sobti, Robert Vandenberg, Alastair G. Stewart, Emad Tajkhorshid, Josep Font, Renae Ryan

**Affiliations:** Transporter Biology Group, School of Medical Sciences, Faculty of Medicine and Health, University of Sydney, NSW, Australia; NIH Center for Macromolecular Modeling and Bioinformatics, Beckman Institute for Advanced Science and Technology, Department of Biochemistry, and Center for Biophysics and Quantitative Biology, University of Illinois at Urbana-Champaign, Urbana, IL 61801, USA; Molecular, Structural and Computational Biology Division, The Victor Chang Cardiac Research Institute, Darlinghurst, NSW 2010, Australia; St Vincent’s Clinical School, Faculty of Medicine, UNSW Sydney, Kensington, NSW 2052, Australia; Department of Physiology and Cellular Biophysics, Columbia University Irving Medical Center, New York, NY, USA

## Abstract

Glutamate is the most abundant excitatory neurotransmitter in the central nervous system, therefore its precise control is vital for maintaining normal brain function and preventing excitotoxicity^1^. Removal of extracellular glutamate is achieved by plasma membrane-bound transporters, which couple glutamate transport to sodium, potassium and pH gradients using an elevator mechanism^2–5^. Glutamate transporters also conduct chloride ions via a channel-like process that is thermodynamically uncoupled from transport^6–8^. However, the molecular mechanisms that allow these dual-function transporters to carry out two seemingly contradictory roles are unknown. Here we report the cryo-electron microscopy structure of a glutamate transporter homologue in an open-channel state, revealing an aqueous cavity that is formed during the transport cycle. Using functional studies and molecular dynamics simulations, we show that this cavity is an aqueous-accessible chloride permeation pathway gated by two hydrophobic regions, and is conserved across mammalian and archaeal glutamate transporters. Our findings provide insight into the mechanism by which glutamate transporters support their dual function and add a crucial piece of information to aid mapping of the complete transport cycle shared by the SLC1A transporter family.

## Introduction

Excitatory amino acid transporters (EAATs) are glutamate transporters that belong to the solute carrier 1A (SLC1A) family. EAATs couple glutamate transport to the co-transport of three sodium (Na^+^) ions and one proton (H^+^) and the counter-transport of one potassium (K^+^) ion^4^. In addition to this coupled transport, binding of substrate and Na^+^ ions to EAATs activates a reversible and thermodynamically uncoupled chloride (Cl^−^) conductance. Mutations that alter the properties of this Cl^−^ conductance have been linked to episodic ataxia type 6 (EA6)^9, 10^.

X-ray crystal structures and, more recently, single-particle cryo-electron microscopy (cryo-EM) structures of several members of the SLC1A family have revealed distinct conformations within the transport cycle. Structures of EAAT1^11^, neutral amino acid transporter 2 (ASCT2)^12–14^, and the archaeal homologs, Glt_Ph_^2, 15–22^ and Glt_Tk_^23–25^ have shown that SLC1A transporters share a similar trimeric structure in which each protomer, comprising a transport and a scaffold domain, can function independently^25–27^. EAATs utilize a twisting elevator mechanism of transport, in which the transport domain, containing the substrate and co-transport ion binding sites, shuttles across the membrane during transport while the scaffold domain makes inter-subunit contacts and remains anchored to the membrane^2, 13^. This results in sequential transitions between an “outward-facing” state (OFS), an “intermediate outward-facing” state (iOFS), and an “inward-facing” state (IFS). A single gate (HP2) opens on the extracellular side to accept substrate and the cytoplasmic side to release it^2, 13, 20^ (Fig. 1a).

**Fig. 1.**
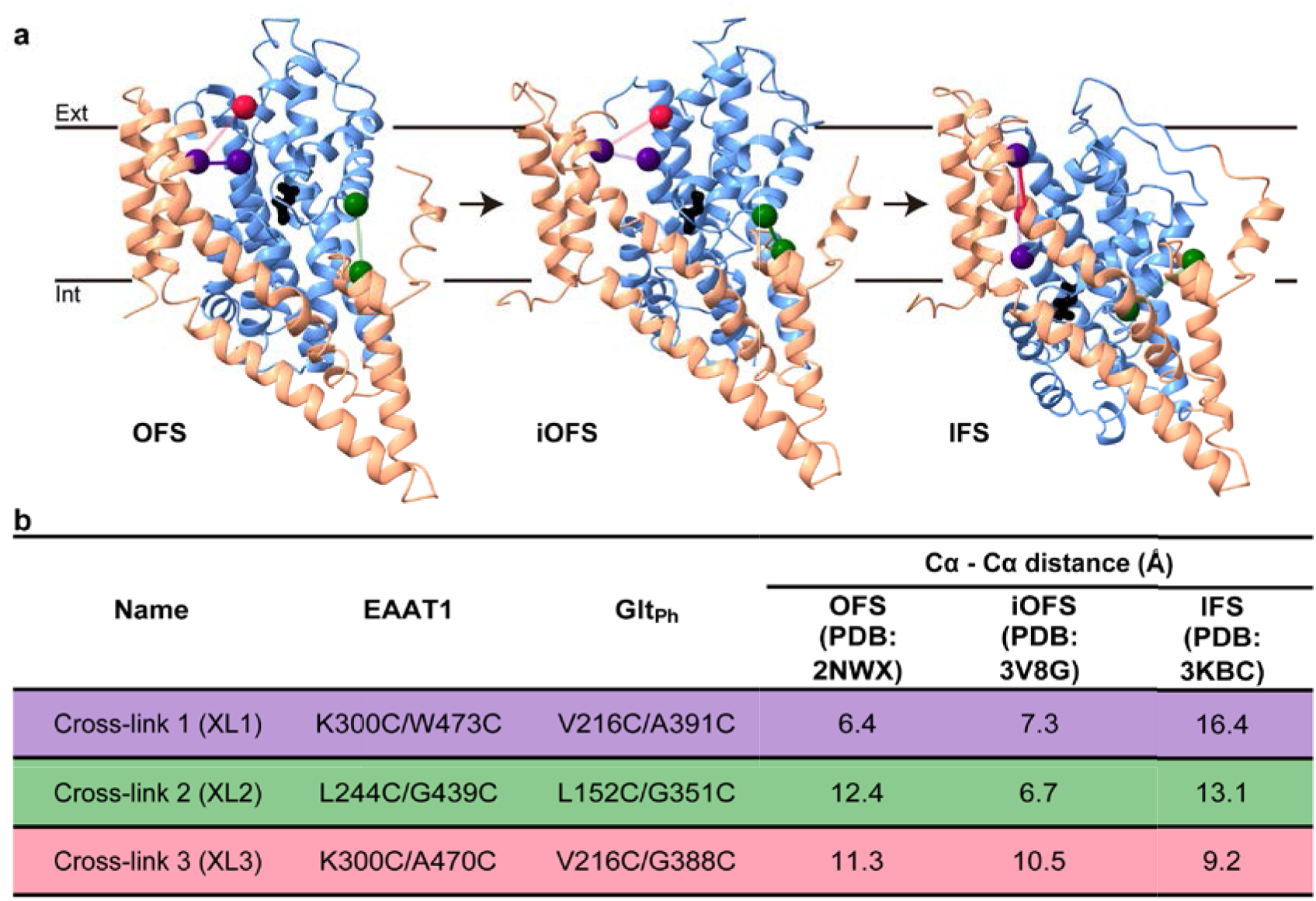
Glt_Ph_ utilizes an elevator mechanism that can be probed by cross-links between the transport and scaffold domains. **a,** Glt_Ph_ protomer viewed parallel to the membrane in the OFS (PDB: 2NWX), iOFS (PDB: 3V8G) and IFS (PDB: 3KBC). The static scaffold domain is shown in salmon, the mobile transport domain in blue and the substrate, aspartate, represented by black sticks. The three double cysteine pairs designed for cross-linking, XL1, XL2 and XL3, are shown in purple, green and pink, respectively. Solid color reflects the approximate conformation in which each cross-link is designed to trap. **b,** Cα-Cα distances of cysteine pairs according to the corresponding residues in the known crystal structures of Glt_Ph_.

A clear pathway for the uncoupled Cl^−^ conductance has not been observed in the available structures. Therefore, the molecular determinants required for activation remain unclear. Electrophysiological studies have identified several residues in EAAT1 that likely line the channel pore based on their ability to alter Cl^−^ conductance properties, and not transport properties, when mutated^7, 28–34^. Some of these residues line an aqueous cavity that was observed at the interface of the transport and scaffold domain in the iOFS, leading to the hypothesis that the cavity may be a partially formed Cl^−^ channel that becomes fully open in a different conformational state^20, 28^.

Here we investigate the twisting elevator mechanism of the transport domain to explore its relationship to the opening of the Cl^−^ channel, and thus, the interplay between the dual functions of EAATs. Using a disulfide cross-linking strategy, we design three cysteine pairs to trap Glt_Ph_ in different states within the transport cycle (Fig. 1a, b), one of which yields a novel structure in an open-channel conformation. We also present molecular dynamics (MD) simulations and functional experiments in Glt_Ph_ and EAAT1, which confirm the cavity at the domain interface is an aqueous-accessible Cl^−^ permeation pathway conserved across mammalian and archaeal SLC1A transporters.

## Results

### Glt_Ph_ can be trapped in an open-channel conformation

To gain further understanding of the transport cycle of glutamate transporters, and to uncover how they enter into an ion-conducting state, we utilized a disulfide cross-linking strategy to isolate different stages of the transport cycle in Glt_Ph_ – an archaeal homolog of EAATs that transports aspartate. Three cysteine pairs were introduced into fully functional cysteine-less Glt_Ph_ constructs^35^ such that one cysteine residue was located on the mobile transport domain and the other on the relatively static scaffold domain (Fig. 1a). The positions of the cysteines in each pair were chosen for their predicted proximity in the OFS (XL1), iOFS (XL2) and IFS (XL3) stages of the transport cycle (Fig. 1b). The homobifunctional cross-linking reagent, mercury (HgCl_2_), was used to promote complete cross-linking within each double cysteine mutant to ensure sample homogeneity and was verified using an SDS-PAGE gel-shift assay^34^ (Extended Data Fig. 1a).

The three constructs, referred to as Glt_Ph_-XL1, XL2 and XL3, were crystallized for structural determination. Glt_Ph_-XL1 and Glt_Ph_-XL3 were solved at a resolution of 3.8 Å and 3.5 Å resolutions, respectively (Extended Data Table 1). The structure of Glt_Ph_-XL1 was similar to the OFS (Cα RMSD = 0.5 Å) and Glt_Ph_-XL3 similar to the IFS (Cα RMSD = 0.6 Å) (Extended Data Fig. 1b). A continuous pore was not observed in either of these structures, in agreement with previous strudies^2, 16^. Despite extensive screening and crystal optimization, the structure of Glt_Ph_-XL2 could not be determined using crystallographic methods. Instead, purified Glt_Ph_-XL2 protein was reconstituted into nanodiscs and examined using cryo-EM. Samples were prepared using physiologically relevant concentrations of NaCl and aspartate. Full cross-linking of all protomers was ensured by the addition of HgCl_2_ and confirmed by SDS-PAGE (Extended Data Fig. 1c). 3D classification was performed by imposing a C3 symmetry, yielding four classes, two of which were similar consisting of a total of 220,938 particles (Extended Data Fig. 2, Extended Data Table 2). Initial rounds of refinement revealed that all three protomers were in a conformation similar to the iOFS crystal structure^20^.

Symmetry expansion was used to identify any structural variation between protomers within the Glt_Ph_-XL2 trimer. Particles representing individual protomers were subject to focused 3D classification, which revealed ten classes representing two distinct entities (Extended Data Fig. 2). Of these, nine classes adopted a conformation resembling the iOFS, whereas one class exhibited considerable differences. The latter, unique class and also the clearest class resembling the iOFS were further refined and post-processed to a resolution of 4.0 Å and 3.7 Å, respectively (Extended Data Fig. 2, and 3, Extended Data Table 2). The iOFS structure was essentially indistinguishable from that previously reported (RMSD = 0.9 Å)^20^ (Extended Data Fig. 1d, 4a, b). However, the unique class revealed a novel structure with substantial conformational differences to the iOFS, including an aqueous-accessible pathway (Fig. 2a, Extended Data Fig. 4a, b). We therefore refer to this novel conformation as the Cl^−^-conducting state (ClCS).

**Fig. 2.**
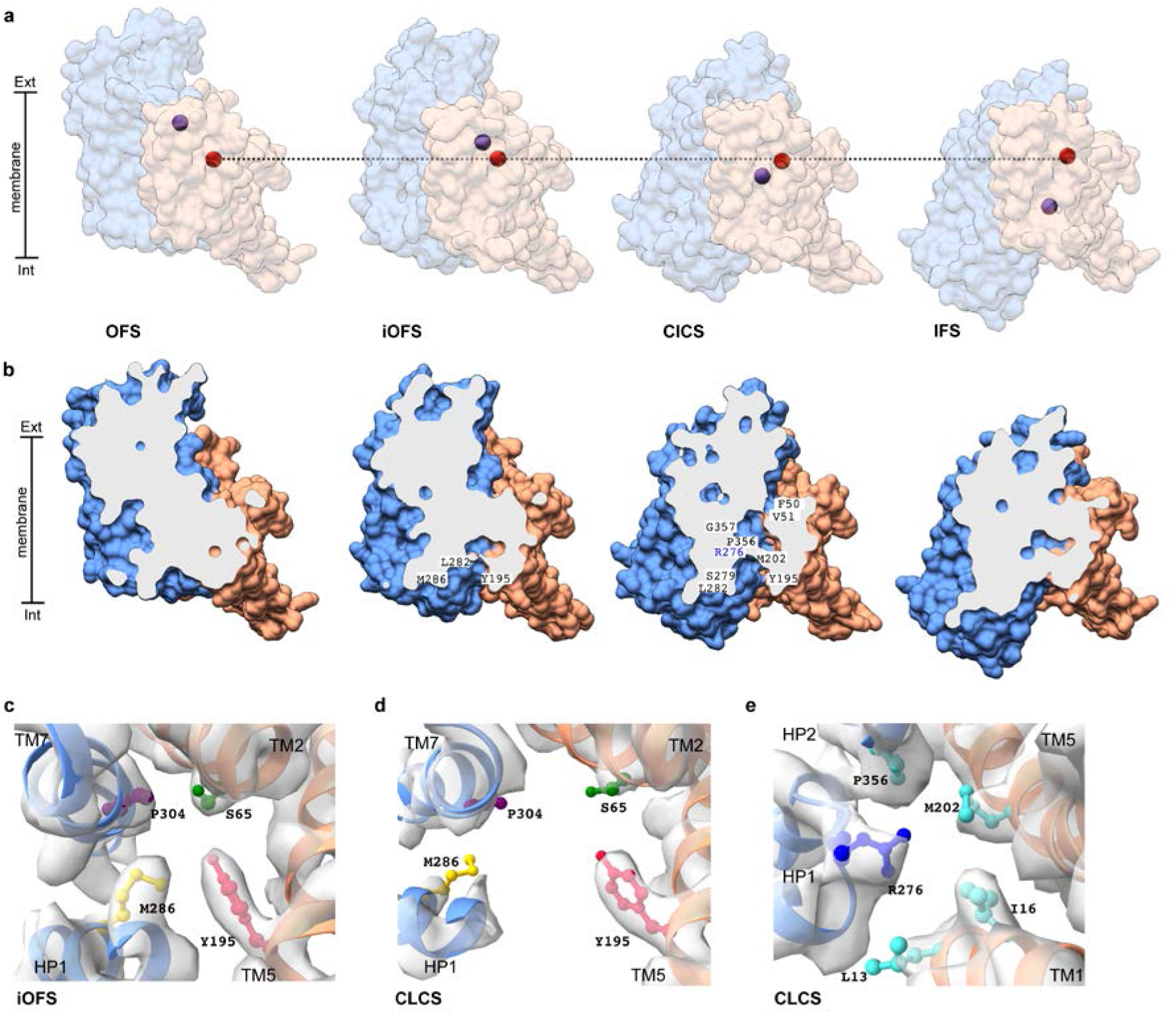
Glt_Ph_ can be trapped in an open-channel conformation. X-ray and cryo-EM structures of Glt_Ph_ in different states. **a,** Surface representation of Glt_Ph_-XL1 in the OFS, Glt_Ph_-XL2 in the iOFS and ClCS, and Glt_Ph_-XL3 in the IFS, viewed from within the membrane plane. The color scheme is the same as in Fig. 1. The Cα positions of L152C and G351C are shown as spheres in red and blue, respectively. The dashed line through L152C highlights the unchanging position of that residue throughout the transport cycle as G351C moves towards the intracellular space. **b,** Cross-sections through R276 of the protomers in **(a)**. Residues lining the domain interface are labelled. **c, d,** Close-up of the “constriction zone” viewed from the extracellular space in the iOFS **(c)** and the ClCS **(d)** fitted in their respective Cryo-EM maps. **e,**Close-up of the narrowest point of the cavity in the ClCS. Noise removed for clarity.

The scaffold domain (TM1, TM2, TM4 and TM5) is relatively similar in the iOFS and ClCS (Cα RMSD = 0.6 Å between the two). However, the transport domain (TM3, TM6, TM7, TM8, HP1 and HP2) is further towards the IFS in the ClCS, following a ~5.6 Å translation and ~17.5° rotation (Extended Data Fig. 4c). In this novel conformation, the position of G351C, located on the tip of HP2, is ~7.6 Å closer to the cytoplasm compared its position in the iOFS (Fig. 2a). Interestingly, the Cα-Cα distances between the two introduced cysteine residues at positions 152 and 351 are similar in the iOFS and the ClCS (6.5 Å and 6.6 Å, respectively), yet the absolute positions of C152 and C351 are significantly different (C351 is above C152 in the iOFS and below it in the ClCS) (Fig. 2a, Extended Data Fig. 4d, e). Although a density for Hg^2+^ was not observed, we infer that the cysteine residues in both structures are cross-linked because the Cα-Cα distances are within the range required for cross-linking by Hg^2+^ and the energetically favorable OFS was not captured. An additional ~5.5 Å downward movement of the transport domain accompanied by a further ~8.7° rotation would be required to reach the IFS, placing this new ClCS approximately half-way between the iOFS and the IFS and two-thirds of the way along the complete substrate translocation pathway (Extended Data Fig. 4c).

Previously, Verdon and Boudker^20^ observed a cavity lined by hydrophobic residues in TM2 and TM5 at the domain interface of the iOFS, which permitted solvent accessibility from the cytoplasmic side. The further twisted and downward position of the transport domain in the ClCS increases the gap between it and the scaffold domain, effectively dismantling the ‘constriction zone’ formed by S65, Y195, M286 and P304 at the intracellular end of the cavity in the iOFS^36^ (Fig. 2b-d). As the transport domain transitions between the iOFS and ClCS, TM7 pulls away from the scaffold domain to extend the cavity and reveal a cluster of hydrophobic residues in TM2 (F50 and V51) and TM5 (L212). These hydrophobic residues are accessible to the extracellular solvent in the ClCS, consistent with previous proposals that they line the Cl^−^ permeation pathway^34, 36^ (Fig. 2b). This results in an aqueous pathway connecting the extracellular and intracellular milieu, which is not a straight route through the domain interface, but rather a pore that bends around the center of the membrane. The narrowest point of the pore occurs at R276 – the proposed anion selectivity filter for the Cl^−^ channel in the SLC1A family^29, 37^. R276, together with a cluster of solvent-exposed hydrophobic residues, L13, I16, M202 and P356, form a restriction point in the aqueous pathway (Fig. 2e). The estimated diameter of the narrowest part of the pore is ~6 Å (calculated by Hole^38^), which is sufficient to accommodate a dehydrated thiocyanate ion^39^ – the largest and most permeant anion carried by EAATs^8, 40^. Thus, our ClCS structure reveals a putative Cl^−^ permeation pathway in Glt_Ph_.

### EAAT1 and Glt_Ph_ can transition into an open-channel conformation

We sought further atomic evidence that the aqueous pathway in the ClCS functions as an ion-permeable pore by exploring the elevator movement of the transport domain between the OFS and IFS using molecular dynamics (MD) simulations. Biasing steered MD (SMD) simulations were performed on the membrane embedded human-EAAT1 (PDB: 5LLU)^11^ to additionally test whether the pathway was conserved across species. Starting from the fully bound OFS (with three Na^+^ ions and aspartate bound) and employing a time-dependent biasing protocol on two collective variables (see methods) representing the translation (ς) and orientation (θ) change of the transport domain (relative to the scaffold domain), transition to the IFS was achieved (Extended Data Fig. 5, Supplementary Video 1). The effectiveness of the biasing protocol was gauged by monitoring the two collective variables (ς, θ) and how they reach their values in the alternate state (Extended Data Fig. 5a). The aim of the SMD simulations was to provide a structural model for the unknown state of EAAT1 (IFS) and a transition pathway to connect the two end states (OFS and IFS; Extended Data Fig. 5d, Supplementary Video 1).

We monitored the formation of water conduction profiles during the entire OFS to IFS transition in order to capture the ClCS. Routes for water permeation were not observed in the two end states, but one of the intermediate conformations revealed a continuous water pathway at the interface of the transport and scaffold domains (Extended Data Fig. 5d). The distance between the Cα atoms of cysteine pair XL2 (L244 and G439) in this water permeating intermediate was ~6 Å, which is within cross-linking distance (Extended Data Fig. 5b). We also observed a continuous water pathway in simulations of our cryo-EM structure of Glt_Ph_-ClCS after embedding into a lipid bilayer (Extended Data Fig. 5e). This demonstrates that both Glt_Ph_ and EAAT1 can enter a similar intermediate state with a continuous water pathway that connects the extracellular and intracellular milieu. The solvent accessible surface area (SASA) values calculated for residues lining the permeation pathway in the ClCS are higher than those in the OFS (Extended Data Fig. 5c), supporting the involvement of the domain interface in the conduction pathway.

A further simulation in an explicit lipid bilayer was performed on the water-conducting intermediate conformation that we obtained for EAAT1 during our biasing simulations. We used distance restraints on the Cα atoms of L244 and G439 to mimic the experimental cross-link and applied an external electric field to study ion permeation through the water-filled pathway (Fig. 3a). Under these conditions, we observed a full Cl^−^ conduction event through EAAT1 in which the ion entered from the intracellular side and travelled the length of the water pathway to the extracellular side (Fig. 3f, Supplementary Video 2). The permeating Cl^−^ ion interacted with several residues lining the domain interface during its translocation, including S103, which plays an important role in anion permeation (Fig. 3b, Extended Data Fig. 5g, Extended Data Table 3). Mutation of S103 to valine, which significantly reduces permeation of anions in both EAAT1 and Glt_Ph_^7, 34^ prevented water permeation through EAAT1 (Extended Data Fig. 5f, Supplementary Videos 3, 4), highlighting the importance of this serine residue for the common Cl^−^ and water pathway at the interface of the transport and scaffold domains. Interestingly, the permeating Cl^−^ ion lost some of its hydration shell (with respect to Cl^−^ in bulk water) during its journey through EAAT1, but never completely dehydrated (Fig. 3c).

**Fig. 3.**
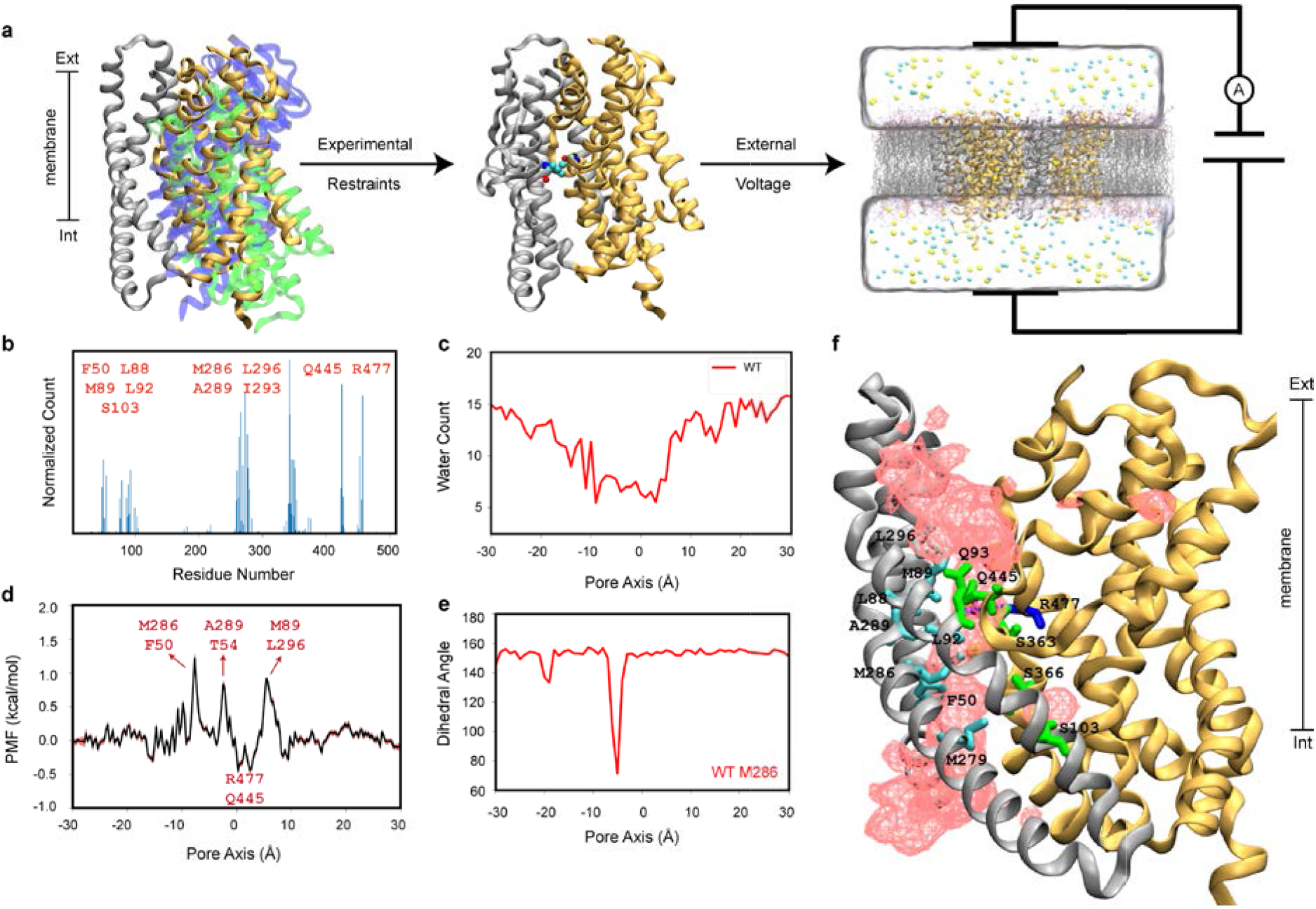
EAAT1 and Glt_Ph_ can transition into an open-channel conformation. **a,** Steered MD simulations of membrane-embedded EAAT1 along translational and orientational degrees of freedom of the transport domain (relative to the scaffold domain) generated an ensemble of structures starting from the OFS (blue) to the IFS (green). The intermediate ClCS (gold) was extracted from the ensemble by measuring the Cα distance between L244-G439, mimicking the experimental cross-link setup. The captured ClCS was equilibrated in an explicit lipid bilayer and subjected to external voltage to drive the movement of Cl^−^. **b,** Residues interacting with the permeating Cl^−^ ion. **c,** The number of water molecules lying within 5 Å of the permeating Cl^−^ ion. **d,** Free-energy of Cl^−^ conduction along the pore axis obtained from umbrella sampling simulations, showing favorable interactions between the permeating Cl^−^ ion and R477/Q445, and barriers against the movement of Cl^−^ formed by two clusters of hydrophobic residues. **e,** The dihedral angle profile of M286 suggesting the re-orientation of this side chain accompanies movement of Cl^−^. **f,** Structure of the EAAT1 protomer showing residues forming the Cl^−^ pathway in green (polar residues), cyan (hydrophobic residues), and blue (basic residues). The occupancy of Cl^−^ is shown in light red isosurface.

We obtained a free-energy profile for Cl^−^ permeation through EAAT1 using umbrella sampling simulations. This revealed two clusters of hydrophobic residues in the extracellular (L88, M89, and L296) and intracellular (F50, T54, M286, and A289) regions of the conduction pathway that provided free-energy barriers against the movement of Cl^−^ (Fig. 3d). Close examination of the simulation trajectories indicated that the free-energy barrier at the intracellular end of the pathway is mainly attributable to the side-chain reorientation of M286 (Fig. 3e). The free-energy profiles also revealed energy minima corresponding to the interaction of permeating Cl^−^ with Q445 (HP2) and R477 (TM8) (Fig 3d; Extended Data Fig. 5h), consistent with the importance of R477 (R276 in Glt_Ph_) for anion selectivity in the Cl^−^ conduction pathway^29, 36, 37^. Together, these data show that both EAAT1 and Glt_Ph_ can transition into a water- and Cl^−^-conducting conformation that resembles our Glt_Ph_ ClCS, and that hydrophobic residues may form a gate at either end of the channel.

### The EAAT1 open-channel conformation conducts Cl^−^

To explore the functional relevance of the different structural states that we isolated in Glt_Ph_ using cysteine pair cross-linking, we introduced the same three cysteine pairs (XL1, XL2 and XL3) into a fully functional cysteine-less EAAT1 (E1) background^41^ and expressed the constructs in *Xenopus laevis* oocytes (Fig. 4). Uptake of L-[^3^H]glutamate increased upon incubation of oocytes with the reducing agent dithiothreitol (DTT), suggesting that all cysteine pairs are able to form a spontaneous disulfide bond. Incubation with the oxidizing agent copper phenanthroline (CuPh), which promotes disulfide formation, resulted in reduced L-[^3^H]glutamate uptake compared to untreated oocytes (Fig. 4a). These effects were only observed when the two cysteines were expressed within an individual protomer (Extended Data Fig. 6a), indicating that the disulfide bond is restricting the elevator movement of the transport domain relative to the scaffold domain and thus inhibiting the transport cycle.

**Fig. 4.**
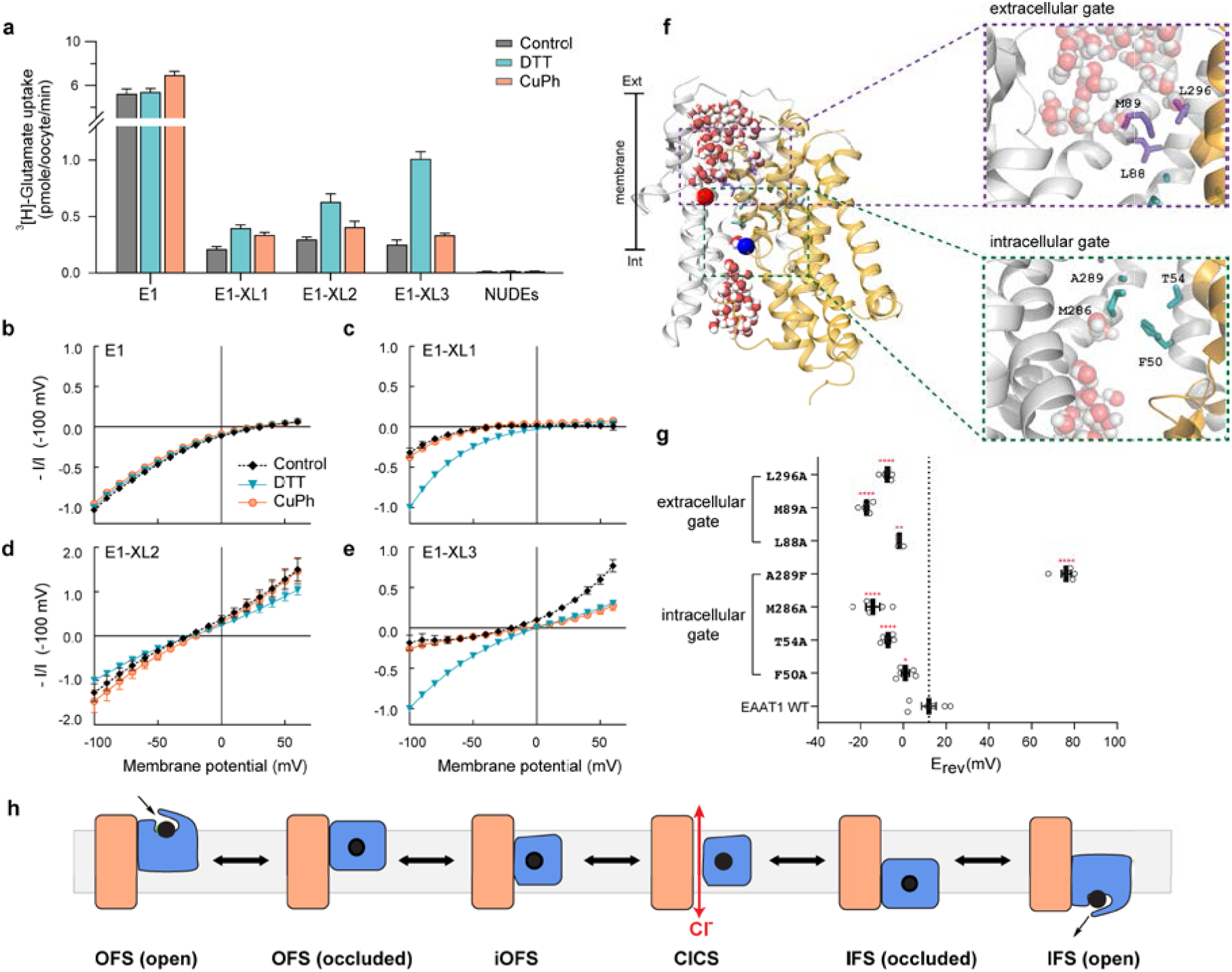
The EAAT1 open-channel conformation conducts Cl^−^. **a,** L-[^3^H]glutamate uptake into oocytes expressing cysteine-less EAAT1 and double cysteine transporter mutants in control conditions (grey), and following pre-incubation with DTT (cyan) or CuPh (orange). **b-e,** L-glutamate elicited current-voltage relationships for cysteine-less E1 **(b)**, E1-XL1 **(c)**, E1-XL2 **(d)** and E1-XL3 **(e)** monitored in the same conditions as **(a)**. Data represent n ≥ 5 ± standard error of the mean (SEM). Note the change of scale in **(d)**. **f,** A snapshot highlighting luminal hydration of EAAT1-IFS with a close-up view of residues forming the extracellular and intracellular hydrophobic gates. The scaffold domain is shown in grey and the transport domain in gold. The Cα atoms of the two introduced cysteine residues are shown as spheres (L244 in red and G439 in blue). **g,** Membrane reversal potentials (E_rev_) measured in oocytes expressing wild-type and mutant EAAT1 transporters. Each white circle represents a response from a single cell. Data represented as mean ± SEM (n ≥ 5). * p < 0.05; ** p < 0.005; **** p < 0.0001. **h,** Schematic representation of the substrate transport cycle. A single protomer is shown with the scaffold domain in salmon, transport domain in blue and substrate in black.

Application of glutamate to oocytes expressing EAAT1 generates an overall conductance that is composed of two components; the coupled transport current and the uncoupled Cl^−^ current (Fig. 4b). The net EAAT1 conductance is inwardly-rectifying and reverses direction at +33 ± 3 mV with the bulk of the current generated at −100 mV due to glutamate transport. In contrast, a linear conductance with a reversal potential of −24 mV is characteristic of the isolated Cl^−^ channel component in *X. laevis* oocytes^6, 42^. Thus, changes in both reversal potential and current amplitude at positive membrane potentials are indicative of alterations to the contribution of Cl^−^ to the total conductance observed. Consistent with the L-^3^[H]glutamate uptake results, incubation of E1-XL1 and E1-XL3 with DTT resulted in a larger glutamate-induced current at −100 mV and incubation with CuPh resulted in a smaller current, revealing a degree of spontaneous cross-linking that inhibits transporter function (Fig. 4c, e). The reversal potential of the E1-XL1 conductance was similar to E1, suggesting that both the transport and channel components were inhibited to the same degree (Fig. 4c). While some outward current observed in E1-XL3 in control conditions, this could not be recovered to the same degree after incubation with CuPh (Fig. 4e). In contrast, the conductance of E1-XL2 was markedly different to E1; the reversal potential was negative (−25.9 ± 2.3 mV) and large currents were apparent at positive membrane potentials under all conditions (Fig. 4d). These features are characteristic of enhanced Cl^−^ channel activity and only observed when the two cysteine residues were within the same protomer (Extended Data Fig. 6b), confirming that an intra-protomer cross-link traps the transporter in a Cl^−^-conducting state. The effects of DTT and CuPh were minimal on the E1-XL2 currents, possibly because of limited accessibility of the introduced cysteine residues to these agents.

To investigate whether the hydrophobic residues that we identified computationally at either end of the permeation pathway have a role in gating the uncoupled Cl^−^ conductance, we mutated each residue to alanine in order to reduce side chain hydrophobicity, except for A289, which was converted to phenylalanine to introduce hydrophobicity (Fig. 4f). When expressed in *X. laevis* oocytes and compared to wild-type EAAT1, the reversal potential of the glutamate-activated conductance formed by the alanine mutants shifted to more hyperpolarized potentials, demonstrating an increase in the proportion of current carried by Cl^−^. This indicated that Cl^−^ can more easily move through the transporter when components of the hydrophobic gates are removed (Fig. 4g). In contrast, A289F resulted in a significant positive shift of the reversal potential to +76 ± 2 mV, virtually eliminating any contribution from Cl^−^ to the glutamate-activated conductance in this mutant. Collectively, these data confirm that the permeation pathway we have identified in EAAT1 forms a functional Cl^−^ channel that is gated by hydrophobic residues at either end.

## Discussion

Glutamate transporters are enigmatic membrane proteins that have an unusual structure. Each of their three protomers forms a twisting elevator that supports a dual transport and channel function. It is thought that the physiological role of the Cl^−^ channel is to reconcile ionic homeostasis and charge balance disrupted by glutamate transport^1, 43^. In the retina, activation of the EAAT5 Cl^−^ channel contributes to a negative feedback loop that leads to a reduction in glutamatergic excitatory transmission^44, 45^. In cerebellar astrocytes, EAAT1/2 channel activity can influence the resting concentration of intracellular Cl^−46^. Disruption of EAAT1 Cl^−^ channel activity underlies the pathogenesis of EA6^9, 10^, a disorder characterized by periods of imbalance and incoordination that can be associated with other neurological symptoms, including seizures and migraine. The P290R mutation results in reduced glutamate transport activity and a large increase in Cl^−^ channel activity, the latter property causing an episodic ataxia-like phenotype and affecting glial cell morphology in a drosophila model^9, 10^.

Using a combination of structural, computational and functional methods, we have successfully mapped the conformational transitions required for dual function glutamate transporters to enter a Cl^−^ conducting state. We have presented the cryo-EM structure of a novel intermediate of the archaeal glutamate transporter homologue, Glt_Ph_, trapped in a Cl^−^ conducting state (ClCS). The structure reveals a continuous aqueous pore at the interface of the scaffold domain and the transport domain, which permits solvent accessibility from either side of the membrane and is lined by residues proposed to line the Cl^−^ permeation pathway^7, 28–34, 36^. Together with structures trapped in the OFS, iOFS and IFS in the transport cycle, our data reveal that the transport domain undergoes a twisting vertical movement as it moves along the translocation pathway from the iOFS to the IFS via the ClCS (Fig. 4h). We have also identified the corresponding ClCS state of EAAT1 using MD simulations and shown that it can conduct water and Cl^−^. Because simulations of both EAAT1 and Glt_Ph_ identify a similar aqueous accessible pathway, we suggest there is a high degree of conservation across SLC1A family members.

Our findings are in contrast to previous studies in Glt_Ph_, including MD simulations suggesting that channel opening results from a lateral movement of the transport domain rather than an intermediate state in the transport cycle, and functional recordings implying that Glt_Ph_ trapped in an outward-facing state (XL1) could enter a Cl^−^-conducting conformation^37, 47^. Indeed, our functional experiments indicate that cross-links trapping the transporter in the OFS inhibit both transport and Cl^−^ channel activities. We do not observe any evidence of a Cl^−^ conducting state in XL1, either in electrophysiological recordings of E1-XL1 or in the crystal structure of Glt_Ph_-XL1. Furthermore, the Cα-Cα distance between V216 and A391 in the ClCS is 14.7 Å, which is outside the range required for cross-linking to occur.

We have also provided further characterization of the molecular determinants of anion permeation and identified two clusters of hydrophobic residues that are present at the intracellular and extracellular regions of the Cl^−^ channel pathway. On the intracellular side of EAAT1, the hydrophobic pocket consists of F50, T54 (TM1), M286, A289 (TM5), whereas L88, M89 (TM2) and L296 (TM5) constitute the hydrophobic cluster on the extracellular side. Our data reveal that these two hydrophobic regions act as gates that allow Cl^−^ ions to enter the pathway from each side of the membrane before encountering the narrowest region near to the arginine residue that is known to determine the anion selectivity of this uncoupled conductance (R477 in EAAT1 and R276 in Glt_Ph_)^29, 37^. Residues that constitute the extracellular gate were previously identified as pore-lining residues using the substituted cysteine accessibility method^36^, however our study is the first to reveal their role as one of two hydrophobic gates. Similar dual hydrophobic gating mechanisms are characteristic of ion channels with a lyotropic selectivity sequence^37^, including calcium-activated Cl^−^ channels^48, 49^ and members of the Cys-loop receptor family^50^. The hydrophobicity of residues that make up the two gates are conserved in other members of the SLC1A family, including the neutral amino acid exchangers ASCT1/2, which also exhibit dual transporter/channel function^51, 52^. It is therefore likely that anion permeation is enabled by common molecular determinants in the entire SLC1A family.

Solvent-accessible hydrophobic residues at either end of the permeation pathway likely render the ClCS an unstable conformation, consistent with predictions that the anion-conducting state occurs with low probability (~0.01) during the transport process^40^. Such a state may be stabilized by the membrane-like environment provided by the nanodiscs used in our study, which could support movement of the transport domain by surrounding it with a compliant lipid bilayer and providing an additional network of protein-lipid interaction^53^. Such membrane perturbations and protein-lipid interactions are enhanced by the introduction of charged arginine residues, which are capable of forming extensive hydrogen bonds with phosphate groups in lipid^54^. Indeed, a proline to arginine mutation in EAAT1 (P290R), which increases the Cl^−^ conductance, and results in EA6, is located near the membrane-water interface in TM5^9, 10^. This suggests that the mutant transporter may remain in an open-channel state for longer due to increased stability of the ClCS, thereby preventing rapid shuttling of the transport domain to deliver its substrate.

In summary, this study has identified the conformational rearrangements and molecular determinants involved in the activation of the Cl^−^ channel of EAAT1, providing a crucial piece of information to map the complete transport cycle shared by the SLC1A transporter family. Our data reveal how the dual transport and channel functions of the glutamate transporters can be achieved in a single membrane protein and may be relevant to understanding other dual function transporters such as the SLC26A9 anion transporter^55^ and neurotransmitter transporters from the SLC6 family^56–59^. Furthermore, this work assists in understanding the functional roles the Cl^−^ channel plays, notably, in maintaining cell excitability and osmotic balance^44, 45, 60^ and provides a framework for the rational development of therapeutics that can differentially modulate substrate transport or channel properties for the treatment of debilitating neurological disorders caused by EAAT dysfunction.

## Methods

All chemicals were purchased from Sigma-Aldrich (Sydney, Australia) unless otherwise stated.

### Site-directed mutagenesis

Cysteine-less EAAT1 (E1) and cysteine-less Glt_Ph_ (CLGlt_Ph_) were subcloned into the oocyte transcription vector (pOTV) and bacterial expression vector (pBAD24), respectively. The pBAD24 vectors encode a 6 x His tag at the C-terminal end of the cysteine-less Glt_Ph_ cDNA. Mutagenesis was performed using the Q5^®^ Site-Directed Mutagenesis Kit (New England BioLabs^®^ Inc.). The cDNA products were purified using the PureLink™ Quick Plasmid Miniprep Kit (Life Technologies) and sequences were verified by the Australian Genome Research Facility (Sydney, Australia).

### Protein expression and purification

CLGlt_Ph_ and double cysteine mutants were transformed into *Escherichia coli (E. coli)* Top10 cells (Invitrogen). Bacterial cells were inoculated in Luria Broth medium containing 100 μg/mL ampicillin at 37 °C until OD_600_ ~0.6-0.8. Protein expression was induced with 0.1% L-arabinose and harvested 4 h post-induction. All protein purification experiments were conducted at 4 °C. Membranes were first isolated and solubilized with 10 mM n-Dodecyl-β-D-maltoside (C_12_M; Anatrace) in binding buffer containing 20 mM HEPES-Tris (pH 7.5), 200 mM NaCl, and 5 mM Na-Asp for 1 h. Proteins were purified using nickel-nitrilotriacetic acid (Ni-NTA) resin (Qiagen) and the histidine tag was subsequently removed by thrombin digestion (10 U/mg of protein). The proteins were further purified by size exclusion chromatography (SEC) using Superdex200 16/60 HiLoad gel filtration column (GE Healthcare). For CLGlt_Ph_, Glt_Ph_-XL1, the column was equilibrated with SEC running buffer containing 20 mM HEPES-Tris (pH 7.5), 25 mM NaCl, 25 mM KCl, 5 mM Na-Asp and 7 mM n-decyl-β-D-maltopyranoside (C_10_M; Anatrace). For Glt_Ph_-XL2 and Glt_Ph_-XL3, the SEC buffer contained 10 mM HEPES-KOH/NaOH (pH 7.5), 100 mM NaCl, 0.1 mM Na-Asp and 7 mM C_10_M. Proteins were concentrated to 6–7 mg/mL for gel analysis and crystallization.

### Mercury-induced cross-linking and SDS-PAGE gel shift assay

Purified proteins were diluted to 1 mg/mL and incubated for 1 h at room temperature in the presence or the absence of 10-fold molar excess HgCl_2_. Both HgCl_2_-treated and untreated protein samples were incubated with 500 μM 5 kDa methoxyl polyethylene glycol-maleimide (mPEG5K) and 0.5% SDS for 2 h at room temperature^61^. All samples were then re-suspended in 12% SDS loading buffer and run on a NuPAGE™ 4–12% Bis-Tris gel (Invitrogen) at 200 V for 45 min. Proteins were visualized via Coomassie blue staining.

### Crystallography, data collection and analysis

Protein solution at 6–7 mg/mL was incubated with 10-fold molar excess HgCl_2_ on ice for 1 h and dialyzed overnight at 4 °C against their respective SEC buffers to remove excess HgCl_2_. All crystals were grown using hanging-drop vapor diffusion. Glt_Ph_-XL1 was crystallized in condition containing 5–23% PEG 1000, 100 mM Li_2_SO_4_, 50 mM citric acid, 50 mM Na_2_HPO_4_ at 4 °C. Glt_Ph_-XL3 was crystallized in condition containing 30–35% PEG 400, 0.2 M MgCl_2_, 0.1 M KCl and 0.025 M Na-citrate pH 3.5–5 at 20 °C. Crystals of Glt_Ph_-XL1 was cryoprotected by soaking in a solution containing 20% PEG 1000, 50 mM citric acid, 50 mM Na_2_HPO_4,_ 100 mM Li_2_SO_4_, 5% glycerol, 2 mM C_10_M followed by solution containing 30% PEG 1000, 50 mM citric acid, 50 mM Na_2_HPO_4,_ 100 mM Li_2_SO_4_, 5% glycerol 2 mM C_10_M. All crystals were flash-frozen in liquid nitrogen. All crystallographic data were collected on the Eiger 16M detector at the Australian Synchrotron (ACRF ANSTO) beamline MX2 at a wavelength of 0.976 Å^62^. All crystal diffraction datasets were indexed, integrated and scaled using XDS^63^ and the CCP4 software package^64^. One crystal per structure was used for analysis. Initial phases were obtained by molecular replacement with PHASER^65^ using published structures of Glt_Ph_ in the OFS (PDB: 2NWX), the IFS (PDB: 3KBC) and the iOFS (PDB: 3V8G) as search models. The protein models were then manually built in COOT^66^ and further refined using REFMAC5^67^ with TLS and three-fold non-crystallographic three-fold NCS restraints^68^. Phases were further improved by rounds of manual rebuilding followed by restrained refinement in REFMAC5 and PHENIX^69^ with tight three-fold NCS restraints. Unit cell parameters, data collection and refinement statistics are presented in Extended Data Table 1. Structural figures and animations were prepared using PyMOL^70^, Chimera^71^ or VMD^72^.

### Cryo-EM sample preparation and imaging

Membrane scaffold protein (MSP1E3) was expressed and purified in *E. coli* BL21(Gold) cells and purified Glt_Ph_ proteins were reconstituted into nanodiscs as described previously^73^. Briefly, *E.coli* total lipid extract and 1-parmitoyl-2-oleoyl-glycero-3-phosphocholine (POPC) (Avanti) dissolved in chloroform were mixed in a ratio of 3:1 and dried under a gentle stream of nitrogen. The dried lipid film was resuspended in buffer containing 10 mM HEPES/KOH pH 7.5, 100 mM NaCl, 0.1 mM Na-Asp and 49 mM C_10_M by freeze-thaw cycles until the mixture was clear, which resulted in 33.5 mM lipid stock. Glt_Ph_ proteins were mixed with MSP1E3 and the lipid stock in a molar ratio of 1:1:50 where the final lipid concentration was 4 mM and then incubated at 4 °C for 30 min. An equal volume of Biobeads SM-2 (Bio-Rad) was added to the nanodisc mixture for detergent removal, incubated at room temperature for 1 h on a rotator and then overnight at 4 °C upon replacing with fresh biobeads. The protein-embedded nanodiscs were isolated using a Superose 6 Increase 10/300 GL column (GE Lifesciences) in buffer containing 10 mM HEPES/KOH pH 7.5, 100 mM NaCl and 0.1 mM Na-Asp and concentrated to 4 mg/mL.

3.5 μL of protein-embedded nanodiscs supplemented with 1.5 mM Fos-Choline-8 (Anatrace) was applied to a glow-discharged holey carbon grid (Quantifoil R1.2/1.3, 200 mesh). Using Vitrobot Mark IV (FEI), grids were blotted for 3 s at 22 °C and 100% humidity, before flash-freezing in liquid ethane.

### Data collection and processing

Grids were imaged with a Talos Arctica transmission electron microscope (Thermo Fisher Scientific) at 200 kV equipped with a Falcon 3EC detector ((Thermo Fisher Scientific)) operating in counting mode. Data collection was performed at 150,000 x magnification, resulting in a physical pixel size of 0.986 Å. Each movie was exposed to a total dose of 40 electrons per Å over 29 frames with a total exposure time of 55 s. A total of 2,419 movie micrographs were collected over two sessions and merged together.

Data processing was performed in RELION 3.1^74^. The frame stacks were aligned and corrected for local beam-induced motion using MotionCorr2^75^ with 5 × 5 patches. Contrast transfer function (CTF) estimation was performed with CTFFIND^76^ and micrographs with poor statistics were discarded, yielding 1,843 micrographs that were used for further analysis. 1,199 particles were manually picked from multiple micrographs and subsequently subjected to 2D classification to create templates for autopicking. 731,422 particles were automatically picked, extracted and subjected to one round of 2D classification. An initial model was generated using 311,380 particles from the best twelve class averages from the 2D classification. 3D classification was performed using the initial model low pass filtered to 50 Å as reference, resulting in four 3D classes. Of those, 220,938 particles from two classes (that exhibited the indistinguishable trimeric assembly of Glt_Ph_ and conformation) were combined, re-extracted and subjected to 3D refinement, post-process and Bayesian polishing. The resulting map of 3.9 Å resolution showed all protomers in a conformation similar to the iOFS (PDB: 3V8G). Symmetry expansion^77^ was performed on 662,814 monomer particles to probe for structural heterogeneity between the trimer protomers. The particles were then subjected to one round of 3D classification without image shifts using a regularization parameter of 40, yielding ten 3D classes. Of the ten 3D classes, nine classes closely resembled the iOFS conformation and one class consisting of 14.8% of the symmetry expanded particles was in the ClC conformation. The best iOFS class and ClC were selected for further refinement with a local mask excluding densities attributed to nanodiscs. The final resolution for the iOFS and the ClC monomers were 3.7 Å and 4.0 Å, respectively, using the gold-standard procedure^78^.

### Model building

One subunit of Glt_Ph_ in the iOFS (PDB: 3V8G, Chain C) and the IFS (PDB: 3KBC) were docked into the density map of the iOFS and the ClC, respectively, using PHENIX^79^. For the trimer, three subunits of the iOFS were docked into the map. All models were refined by several rounds of real-space refinement in PHENIX^80^, and model building in coot^66^ and ISOLDE^81^.

## Computational Methods

### Membrane embedding and equilibrium molecular dynamics (MD) simulations of EAAT1

The simulation system was designed based on the OFS conformation of human EAAT1cryst-II (EAAT1) construct (PDB: 5LLU)^11^. Protonation states of titratable residues were assigned using PROPKA^82^. The Membrane Builder module of CHARMM-GUI^83^ was used to embed EAAT1 in a pre-equilibrated patch of membrane containing 1-palmitoyl-2-oleoyl-sn-glycero-3-phosphocholine (POPC) and cholesterol (Chol) in the ratio of 1:1. The system was solvated and neutralized with 150 mM of NaCl. The final constructed system contained ~220,000 atoms with dimensions of 150 × 150 × 97 Å^3^ prior to any simulations.

All MD simulations were performed using NAMD2^84^ with the CHARMM36 protein and lipid force fields^85^ and TIP3P water. The simulations were performed at a temperature of 310 K which was maintained using Langevin thermostat with a damping coefficient of γ=1.0 ps-1, and the Nosé-Hoover Langevin piston method^86^ was used to maintain constant pressure (1 bar). All long-range electrostatic forces were calculated using the particle mesh Ewald (PME) method^87^, and a 2 fs time step was used for integration. Non-bonded interactions were calculated with a cutoff of 12 Å and a switching distance of 10 Å. The system was initially energy minimized and equilibrated for 5 ns with protein backbone atoms harmonically restrained to their initial positions with a force constant of *k* = 1 kcal/mol/Å^2^. The restraints were released during the production run. All simulations were performed in the fully-bound state of the transporter, i.e. in the presence of the substrate, aspartate and three Na^+^ ions^16^. The unresolved Na^+^ ion (Na3) was modeled based on previous structural and simulation studies^88^.

### Sampling structural transitions from OFS to IFS and identification of Cl^−^ conducting intermediate in EAAT1

Comparison of the EAAT1 structure in the OFS (PDB:5LLU)^11^ and the inward-occluded crystal structure of Glt_Ph_ (PDB:4X2S)^15^, a close analog of EAAT1, indicates that the complete structural transition involves a combination of a translation of the transport domain along the membrane normal (ς) and an orientation change with respect to the scaffold domain (θ). Thus, we employed biasing steered MD (SMD)^89^ simulations with translation and orientation as reaction coordinates (also referred to as collective variables, CVs). Cα atoms of the scaffold domain (residues 50 - 110 and 191 - 281) and the transport domain (residues 113 - 147 and 291 – 439), were used to define the CVs. Harmonic force constants of 10^4^ kcal/mol/rad^2^ and 10 kcal/mol/Å^2^ were used for the orientation-based CVs and distance-based CVs, respectively. 50-ns-long biased simulations were performed to drive the structural transition of EAAT1. The effectiveness of the biasing protocol in producing the target alternate conformational state was gauged by measuring the two CVs and comparing them to the target values.

To capture the ClC from the ensemble of structures generated during the biased simulations of EAAT1, we monitored the formation of the water permeation profiles connecting the two sides of the membrane. Interestingly, water wires are captured in an intermediate protein conformation arising during the transition between the two end states with, a ~6 Å distance between the Cα atoms of the residues L244 and G439, the same residue pair cross-linked experimentally in XL-2. It is to be noted that there can be maximum Cα distance of ~7 Å between cross-linked residues^37^. The obtained conformation was then simulated for 100 ns with distance-based restraints on the Cα atoms of L244 and G439, mimicking the experimental cross-link setup.

### Sampling chloride permeation pathway in EAAT1

To investigate the permeation of Cl^−^ through the intermediate conformation identified from the integrative approach of combining biasing simulations with cross-linking experiments, MD simulations were extended from the fully-equilibrated intermediate conformation described above with varying membrane potentials generated by the application of external electric fields across the simulation system. A uniform electric field was applied perpendicular to the membrane plane in the negative z direction (extracellular to intracellular). Two simulations with transmembrane potentials of 800 mV, and 1 V were performed. Each electric field simulation lasted for 100 ns, during which Cl^−^ movement through the intermediate conformation was monitored. To prevent the disruption of the lipid bilayer under the presence of these high external electric fields, weak harmonic restraints were applied to the phosphorous atoms of the lipid bilayer.

### Free-energy calculations

In order to better sample the Cl^−^ conducting pathway and calculate the associated free-energy cost with the process, we performed umbrella sampling (US) simulations^90^, seeded on the pathway captured from the simulations under the external electric field. US calculations were performed with distance (Z-component) between the Cl^−^ ion and the membrane midplane as the CV. In total 60 windows were generated at a spacing of 1 Å, each simulated for 20 ns with accumulated sampling of 1.2 μs. It is to be noted that US simulations were performed in the fully-bound state with experimentally based distance restraints between the Cα atoms of L244 and G439. All analyses were performed on the last 10 ns trajectory of each window. Free-energy profiles were calculated using the WHAM implementation in LOOS^91^.

### Simulation of the ClC conformation of GltPh obtained using cryo-EM

In order to characterize water permeation through the ClC structure of Glt_Ph_ obtained with cryo-EM, the structure was embedded in a pre-equilibrated patch of membrane containing 1-palmitoyl-2-oleoyl-sn-glycero-3-phosphoethanolamine (POPE), 1-palmitoyl-2-Oleoyl-sn-glycero-3-phosphoglycerol (POPG), and POPC lipids in the ratio of 2:1:1. The unresolved Na^+^ ions were modeled based on the previous studies^16, 88^. The entire system was equilibrated by performing a 100 ns-long MD simulation. A uniform external electric field was applied across the simulation system to facilitate the permeation of water. Two simulations with transmembrane potentials of 800 mV, and 1 V were performed.

### Harvesting and preparing oocytes

Female *Xenopus laevis* frogs were obtained from NASCO (Wisconsin, USA) and harvested as described previously^92^. Briefly, stage V oocytes were harvested following anaesthesia with 6.5 mM tricaine in 7.14 mM sodium bicarbonate, pH 7.5 and stored in OR-2 buffer (82.5 mM NaCl, 2 mM KCl, 1 mM MgCl_2_ 5 mM hemosodium HEPES, pH 7.5). All surgical procedures have been approved by the University of Sydney Animal Ethics under the *Australian Code of Practise for the Care and Use of Animals for Scientific Purposes.* Oocytes were defolliculated by agitation with 2 mg/mL collagenase for 1 h. Following digestion, oocytes were injected with 4.6 ng of cRNA and stored with shaking at 16-18°C in standard frog Ringer’s solution (96 mM NaCl, 2 mM KCl, 1 mM MgCl_2_, 1.8 mM CaCl_2_, 5 mM hemisodium HEPES, pH 7.5) supplemented with 50 μg/mL gentamycin, 50 μg/mL tetracycline, 2.5 mM sodium pyruvate and 0.5 mM theophylline.

### Radiolabeled uptake in oocytes

Uptake of L-[^3^H]glutamate (PerkinElmer Life Sciences) was measured in oocytes expressing E1, double cysteine derivatives and in uninjected oocytes (nudes). Oocytes were incubated in 10 μM L-[^3^H]glutamate for 10 min and uptake was terminated by washing oocytes three times in ice-cold frog Ringer’s solution. Oocytes were then lysed in 1 M NaOH and 1% SDS prior to adding scintillant (Optiphase HisSafe, PerkinElmer). Uptake of L-[^3^H]glutamate was measured using MicroBeta TriLux scintillation counter (PerkinElmer). For cross-linking studies, oocytes were treated with either 1 mM dithiothreitol (DTT) for 1 min or 400 μM copper phenanthroline for 5 min and washed three times in Ringer’s solution at room temperature prior to incubation with 10 μM L-[^3^H]glutamate.

### Electrophysiology

2-4 days after injections, currents were recorded using the two-electrode voltage clamp technique with a Geneclamp 500 amplifier (Axon Instruments, Foster City, CA, USA) interfaced with a PowerLab 2/20 chart recorder (ADInstruments, Sydney, Australia) and a Digidata 1322A (Axon Instruments, CA, USA), which was used in conjunction with Chart software (ADinstuments; Axon instruments). All recordings were made with a bath grounded via a 3 M KCl/ 1% agar bridge linking to a 3 M KCl reservoir containing a Ag/AgCl_2_ ground electrode to minimize offset potentials. Current-voltage relationships for substrate-elicited conductance were determined by measuring substrate-elicited currents during 245 ms voltage pulses between −100 mV and +60 mV at 10 mV steps. Background currents were eliminated by subtracting currents in the absence of substrate from substrate-elicited currents at corresponding membrane potentials. For cross-linking studies, current elicited by an initial control dose of 100 μM L-glutamate was recorded prior to 2 min incubation with 1 mM DTT. Currents were recorded following 1 mM DTT treatment and following subsequent 5 min incubation with 400 μM CuPh. For hydrophobic gate mutagenesis studies, current elicited was measured in Ringer’s solution containing 30 μM aspartate.

### Data availability

All relevant crystallography and cryo-EM data are available from the corresponding author upon request. The maps and the coordinates of the refined models have been deposited into the Protein Data Bank (For Glt_Ph_-XL1, PDB 6X01. Glt_Ph_-XL3, PDB 6WZB. Glt_Ph_-XL2 in the iOFS, PDB 6WYJ, EMDB-21966. For Glt_Ph_-XL2 in the CIC, PDB 6WYK, EMDB-21967. For Glt_Ph_-XL2 trimer in the iOFS, PDB 6WYL, EMDB-21968).

## Extended Data Figures

**Extended Data Fig. 1.**
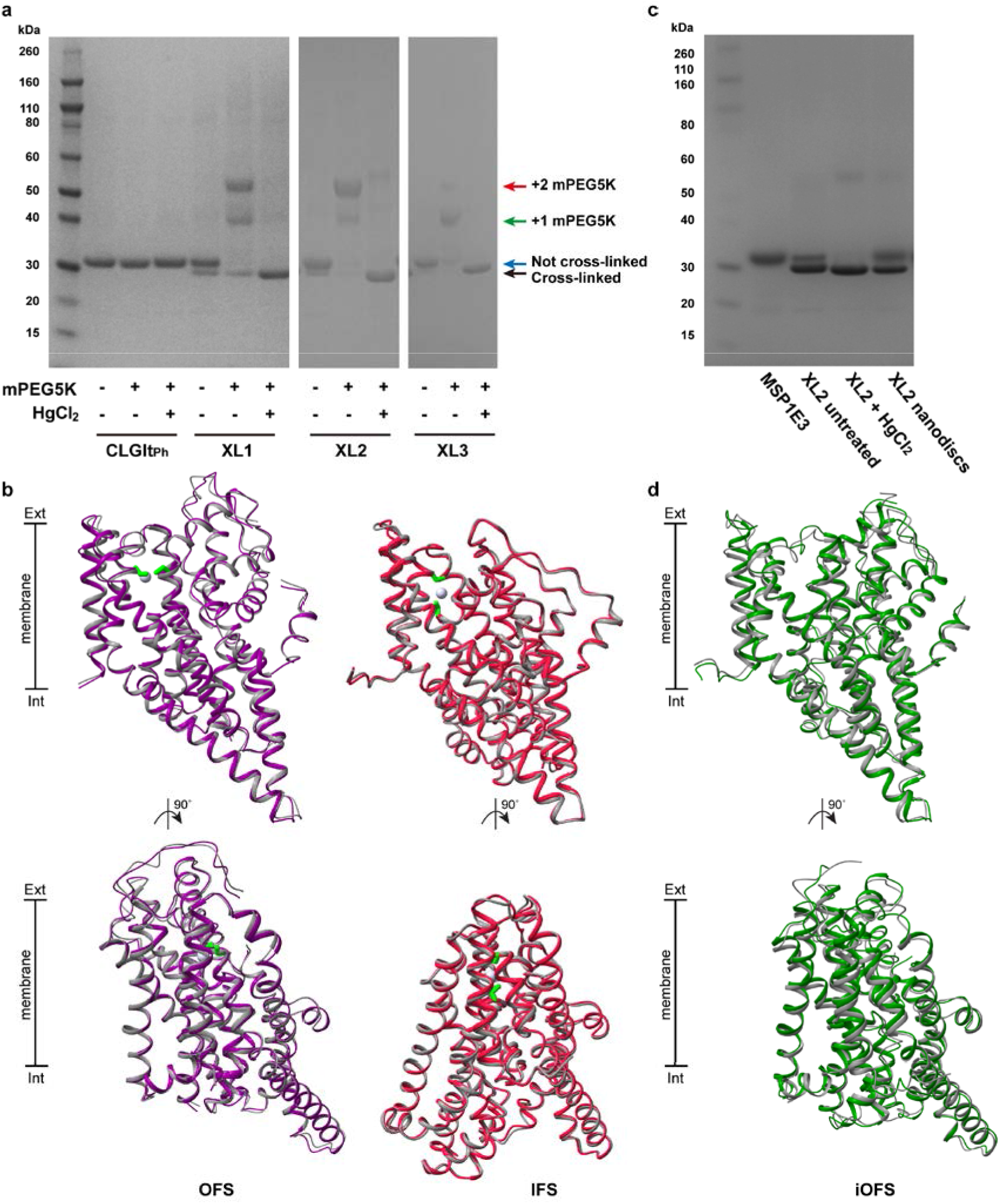
Cross-linking experiments on purified Glt_Ph_ double cysteine mutants. **a,** SDS-PAGE gel shift assay shows the extent of cross-linking in detergent-solubilized mutant Glt_Ph_ under untreated conditions, upon mPEG5K-maleimide treatment and following incubation with HgCl_2_ with arrows indicating the positions of differentially cross-linked protomers and mPEG5K-bound proteins. **b,** Crystal structure of Glt_Ph_-XL1 (purple) and Glt_Ph_-XL3 (pink) superimposed on the OFS (PDB: 2NWX; left, grey) and the IFS (PDB; right, grey), respectively. **c,** SDS-PAGE analysis of purified Glt_Ph_-XL3 in nanodiscs. **d,** Cryo-EM structure of Glt_Ph_-XL2 (green) superimposed on the iOFS (PDB:3V8G, chain C; grey).

**Extended Data Fig. 2.**
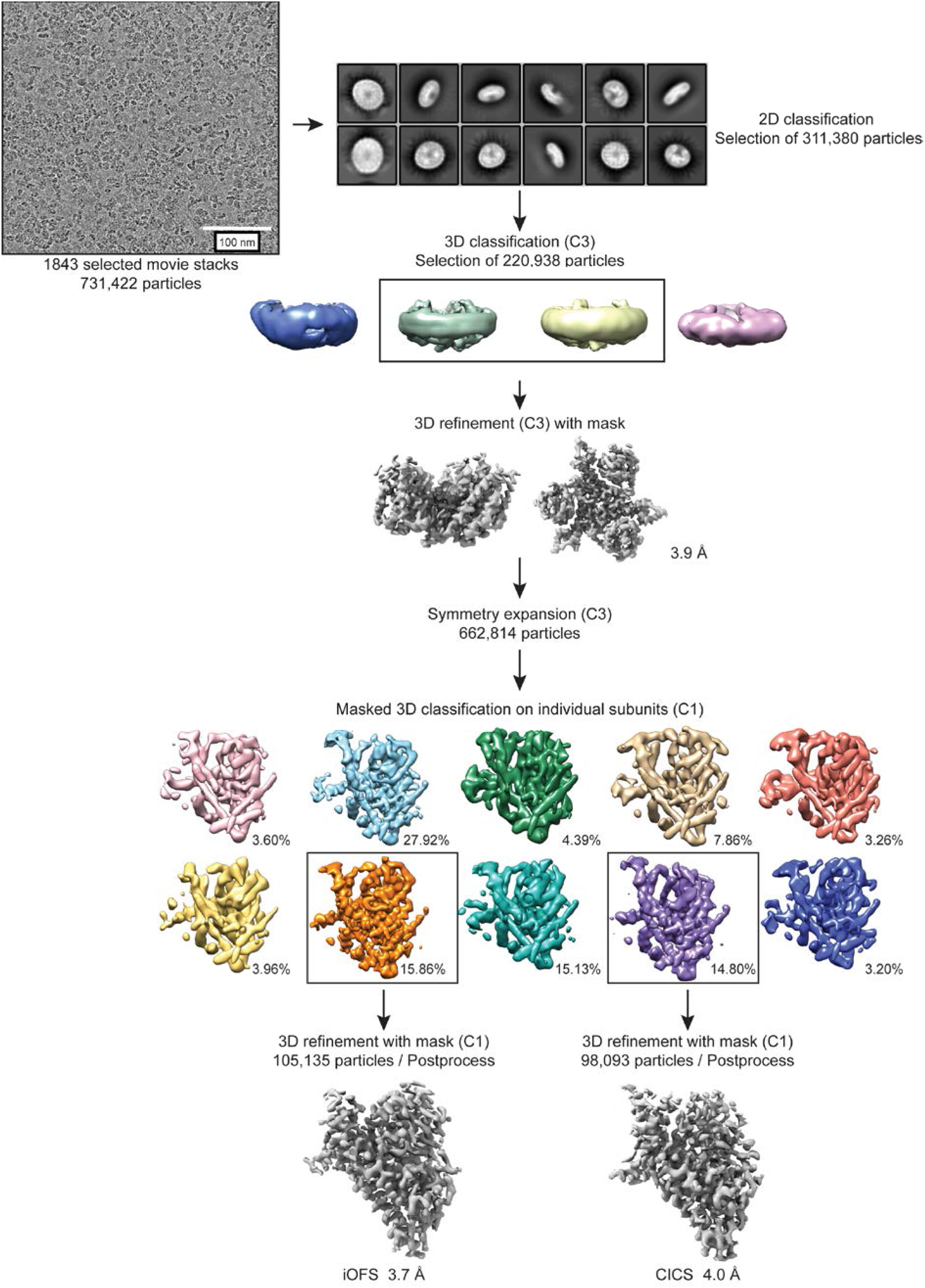
Cryo-EM data processing protocol. Data processing flow chart for Glt_Ph_ reconstituted into nanodiscs in the presence of NaCl and aspartate.

**Extended Data Fig. 3.**
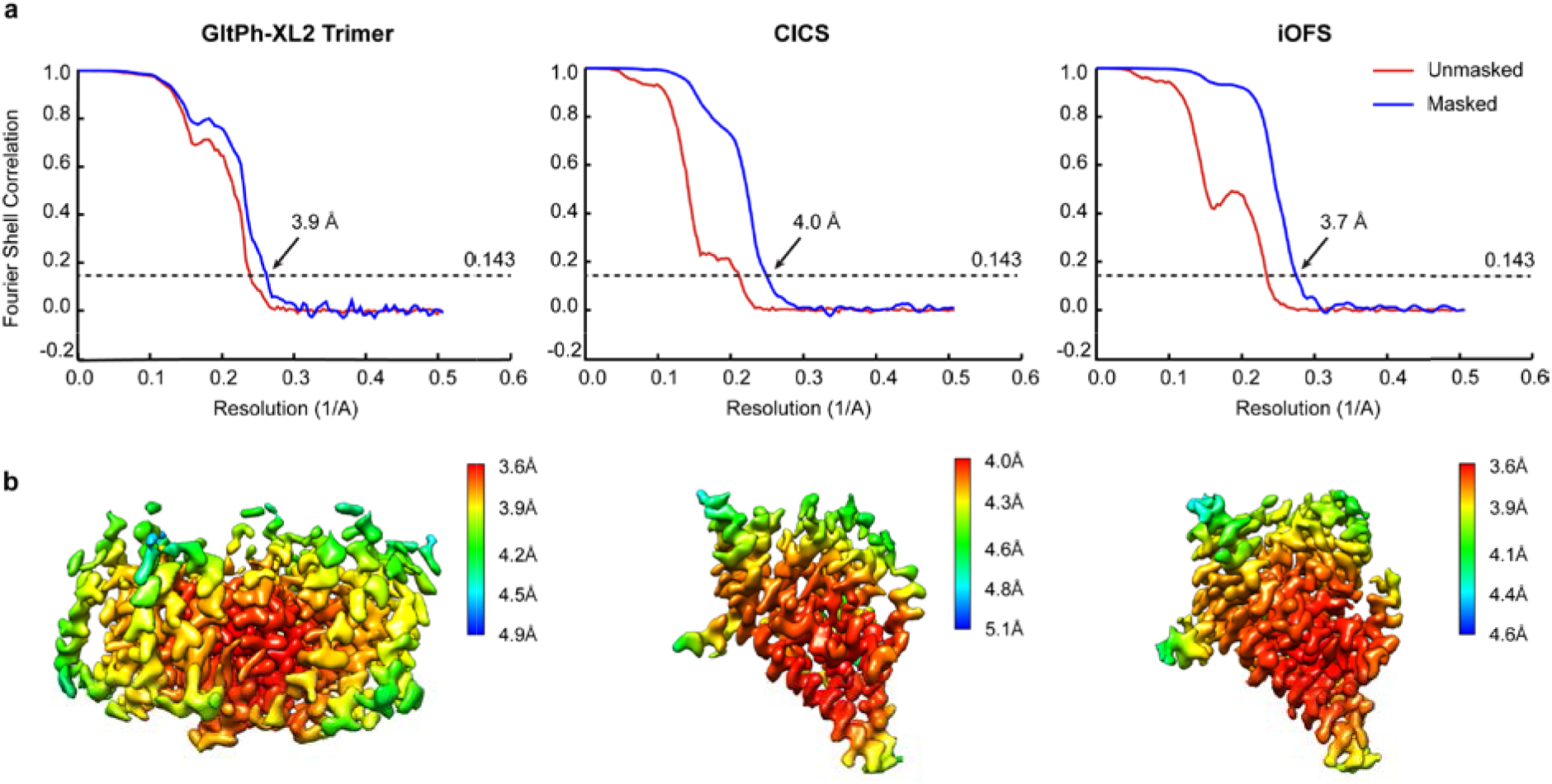
Cryo-EM data refinement. **a,** Fourier shell correlation (FSC) curves indicating the resolution at the 0.143 threshold of final masked (blue) and unmasked (red) maps for Glt_Ph_ trimer iOFS (left), Glt_Ph_ protomer ClCS (middle) and Glt_Ph_ protomer iOFS (right). **b,** Final maps after Relion post-processing, colored according to local resolution estimation using Relion for Glt_Ph_ trimer iOFS (left), Glt_Ph_ protomer ClCS (middle) and Glt_Ph_ protomer iOFS (right).

**Extended Data Fig. 4.**
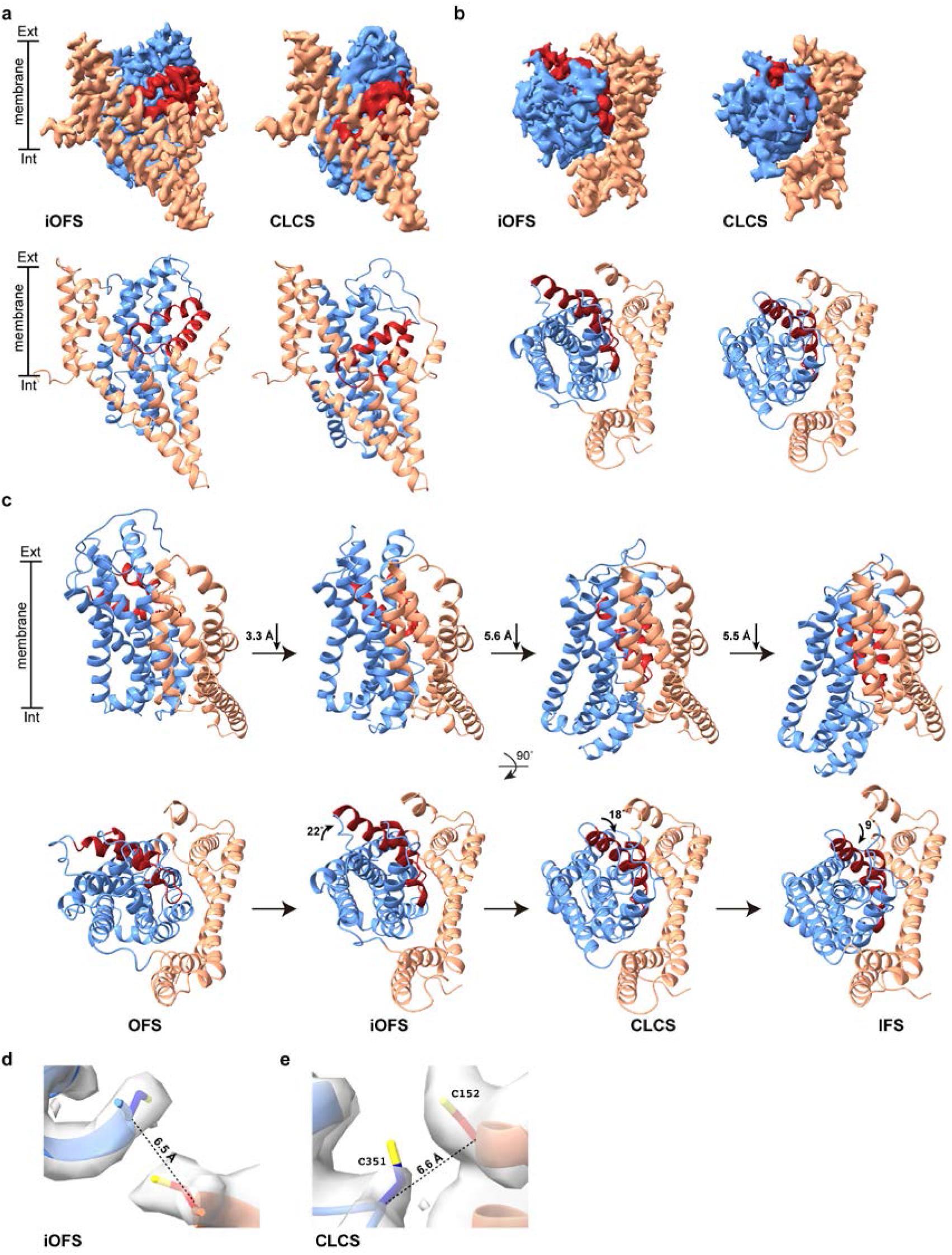
The conformational space of a GltPh protomer. **a,** The front and **b,** top views of the cryo-EM map and atomic model of Glt_Ph_-XL2 in the iOFS and ClCS. Density attributed to the scaffold domain, transport domain and HP2 are shown in salmon, blue and red, respectively. **c,** Conformational changes undertaken by a Glt_Ph_ protomer during the substrate transport cycle viewed from the side and top. HP2 is colored for easier visualization of rotational changes observed in the transport domain. **d, e,** Close-up views of the L152C-G351C cross-link fitted in the iOFS and ClCS cryo-EM maps.

**Extended Data Fig. 5.**
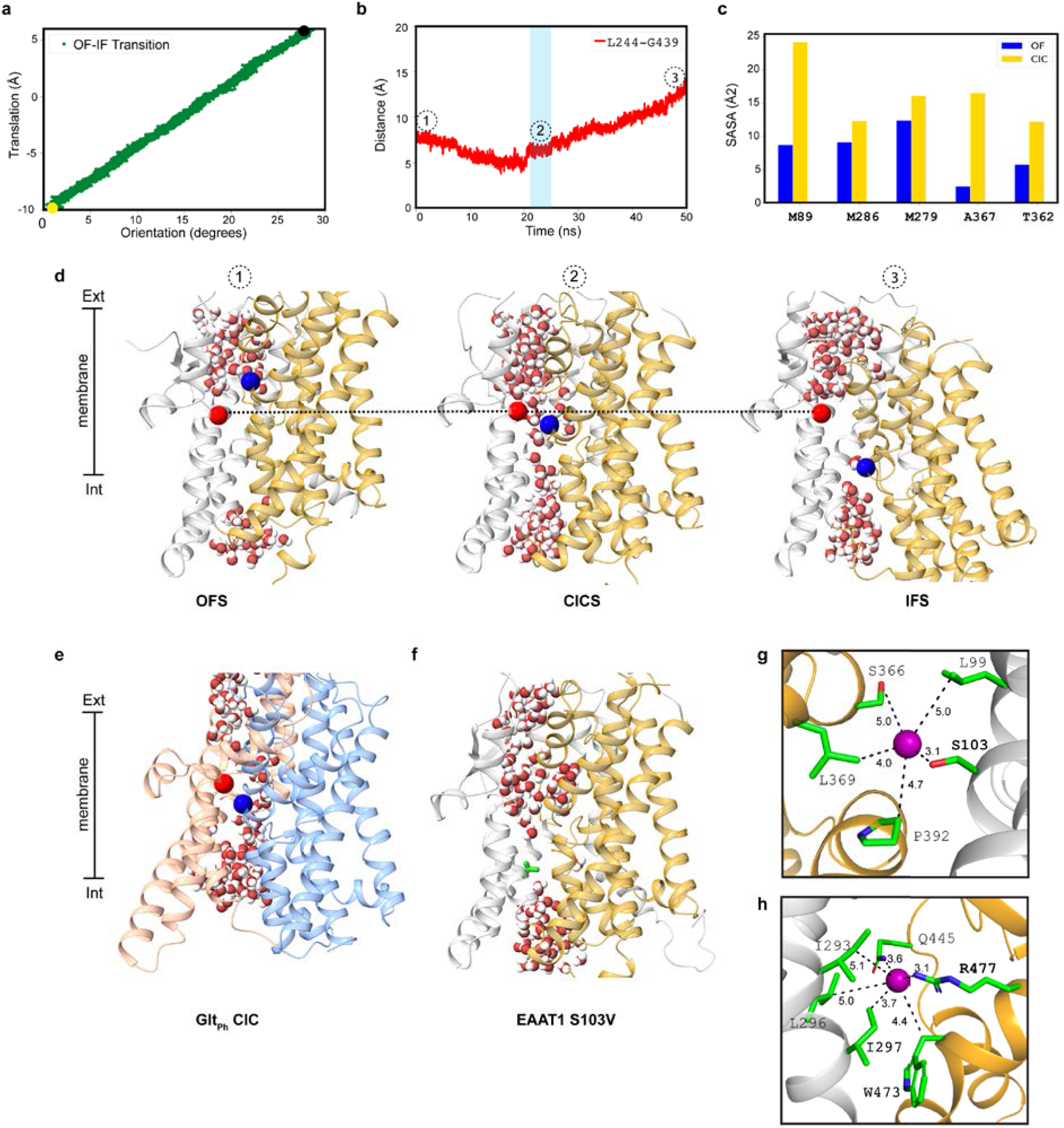
Biasing SMD simulations and water conduction in EAAT1. **a,** Biasing steered MD simulation along translation and orientation degrees of freedom (also referred to as collective variable or CVs) of the transport domain moving with respect to the scaffold domain. Starting from the OFS (black circle), this biasing protocol was used to generate the structurally unknown IFS (yellow circle). **b,** The Cα distance between L244 and G439 (E1-XL2) was monitored over the structural transition between the two end states. Starting from the OFS (1), an intermediate water-conducting state of EAAT1 was captured (2). Blue shading indicates trajectory frames with water permeation pathways connecting the extracellular and intracellular bulk solutions. No water pathway was observed in the OFS (1) or the IFS (3). **c,** Residues lining the Cl^−^ pathway have a higher solvent accessible surface area (SASA) in the ClCS than in the OFS. **d,** Snapshots highlighting luminal water occupancy of the EAAT1 OFS, ClCS and IFS. All biased simulations were performed in fully-bound systems (bound to one aspartate and three Na^+^ ions). The scaffold domain is shown in grey and the transport domain in gold. The Cα atoms of the two introduced cysteine residues are shown as spheres (L244 in red and G439 in blue). **e,** The Glt_Ph_-ClCS structure was embedded into a lipid bilayer containing PE, PG, and PC lipids (mimicking experimental conditions). After an initial equilibration of 100 ns, the entire system was subjected to an external electric field and a continuous water pathway was observed. **f,** When S103 was mutated to valine in EAAT1-ClCS, water molecules were not able to pass through the transporter along the same pathway. Close-up of the region around **g,** S103 and **h,** R477 reveal residues important for Cl^−^ interactions (residues within 5 Å of the Cl^−^ ion labelled).

**Extended Data Fig. 6.**
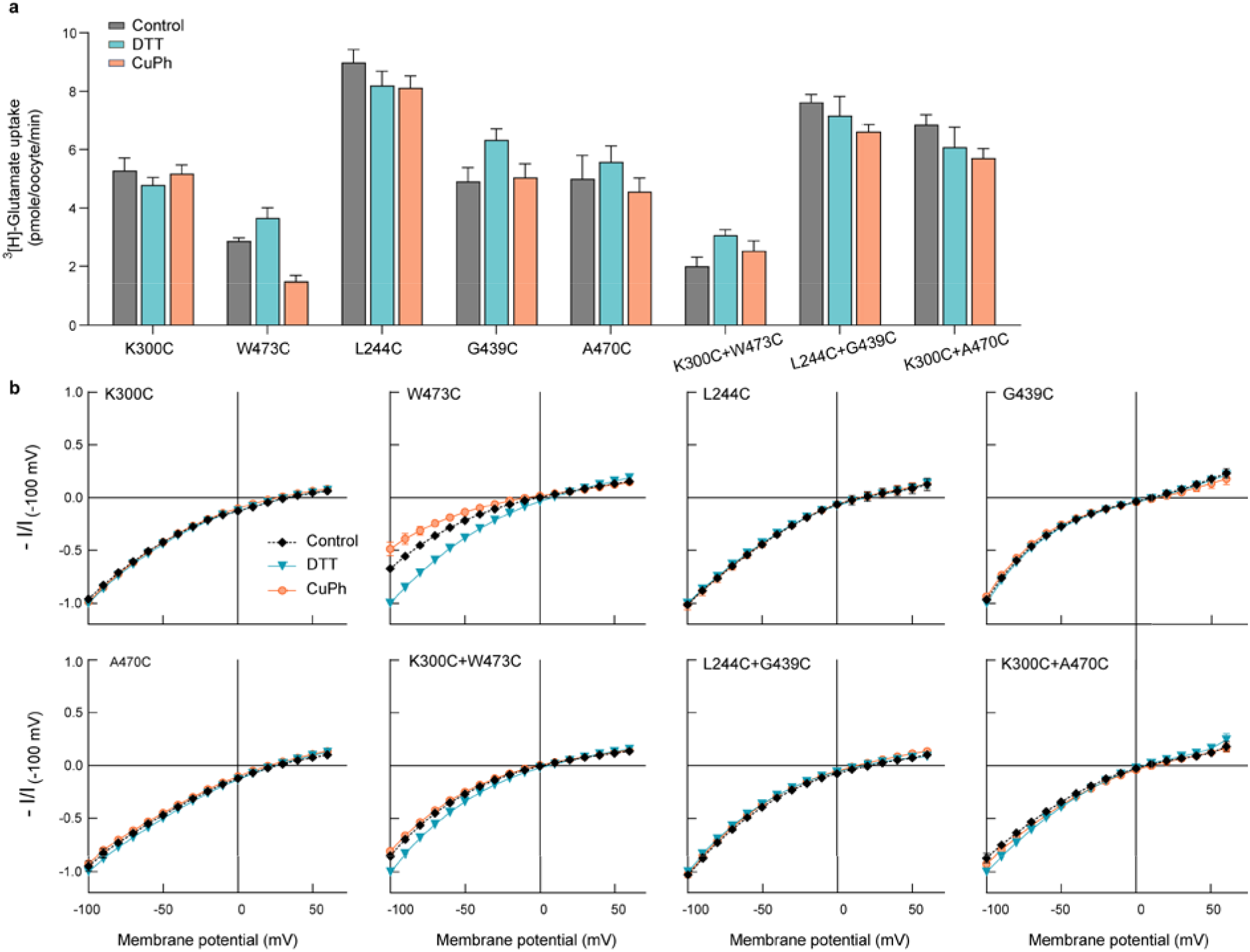
Effect of DTT and CuPh on the single cysteine transporters and co-expressed oocytes. To confirm cross-links E1-XL1, E1-XL2 and E1-XL3 were occurring within an individual protomer rather than between protomers of the trimeric complex, oocytes expressing single cysteine residues either alone or co-injected into an individual oocyte were examined. **a,** L-[^3^H]glutamate uptake under unmodified control conditions (grey), or following pre-incubation with DTT (cyan) or CuPh (orange). **b,** L-glutamate elicited current-voltage relationships for oocytes expressing each single cysteine or co-injected with two single cysteine residues that make up E1-XL1 (K300C and W473C), E1-XL2 (L244C and G439C), and E1-XL3 (K300C and A470C) monitored under the same conditions as **(a)**. Data represent n ≥ 5 ± standard error of the mean (SEM).

**Extended Data Table 1.** Data collection and refinement statistics.

|  | Glt<sub>ph</sub>-XL1<br>(PDB 6X01) | Glt<sub>ph</sub>-XL3<br>(PDB 6WZB) |
| --- | --- | --- |
| **Data collection** |  |  |
| Space group | P6<sub>1</sub> | C222<sub>1</sub> |
| Cell dimensions |  |  |
| *a*, *b*, *c* (Å) | 113.99, 113.99,<br>321.70 | 112.11, 204.57,<br>207.37 |
| $\alpha$, $\beta$, $\gamma$ (°) | 90.00, 90.00,<br>120.00 | 90.00, 90.00,<br>90.00 |
| Resolution (Å) | 48.79 – 3.80 | 49.31 – 3.45 |
| *R*<sub>sym</sub> or *R*<sub>merge</sub> | 0.13 (2.07) | 0.15 (1.52) |
| *I* / σ*I* | 14.00 (2.10) | 10.30 (2.20) |
| Completeness (%) | 100 (100) | 100 (100) |
| Redundancy | 21.70 (21.70) | 13.80 (14.30) |
| **Refinement** |  |  |
| Resolution (Å) | 44.84 – 3.8 | 49.31 – 3.45 |
| No. reflections | 23152 | 31734 |
| *R*<sub>work</sub> / *R*<sub>free</sub> | 25.57/29.58 | 23.46/26.70 |
| No. atoms | 8831 | 9144 |
| Protein | 8822 | 9135 |
| Ligand/ion | 12 | 12 |
| Water | N/A | N/A |
| *B*-factors | 190.88 | 111.47 |
| Protein | 190.85 | 111.45 |
| Ligand/ion | 220.70 | 130.31 |
| Water | N/A | N/A |
| R.m.s. deviations |  |  |
| Bond lengths (Å) | 0.002 | 0.003 |
| Bond angles (°) | 0.50 | 0.62 |
| Ramachandran (%) |  |  |
| Favored | 94.61 | 96.66 |
| Allowed | 4.38 | 3.34 |
| Disallowed | 1.01 | 0.00 |

One crystal per structure was used for analysis.
Values in parentheses are for highest-resolution shell.

**Extended Data Table 2.**
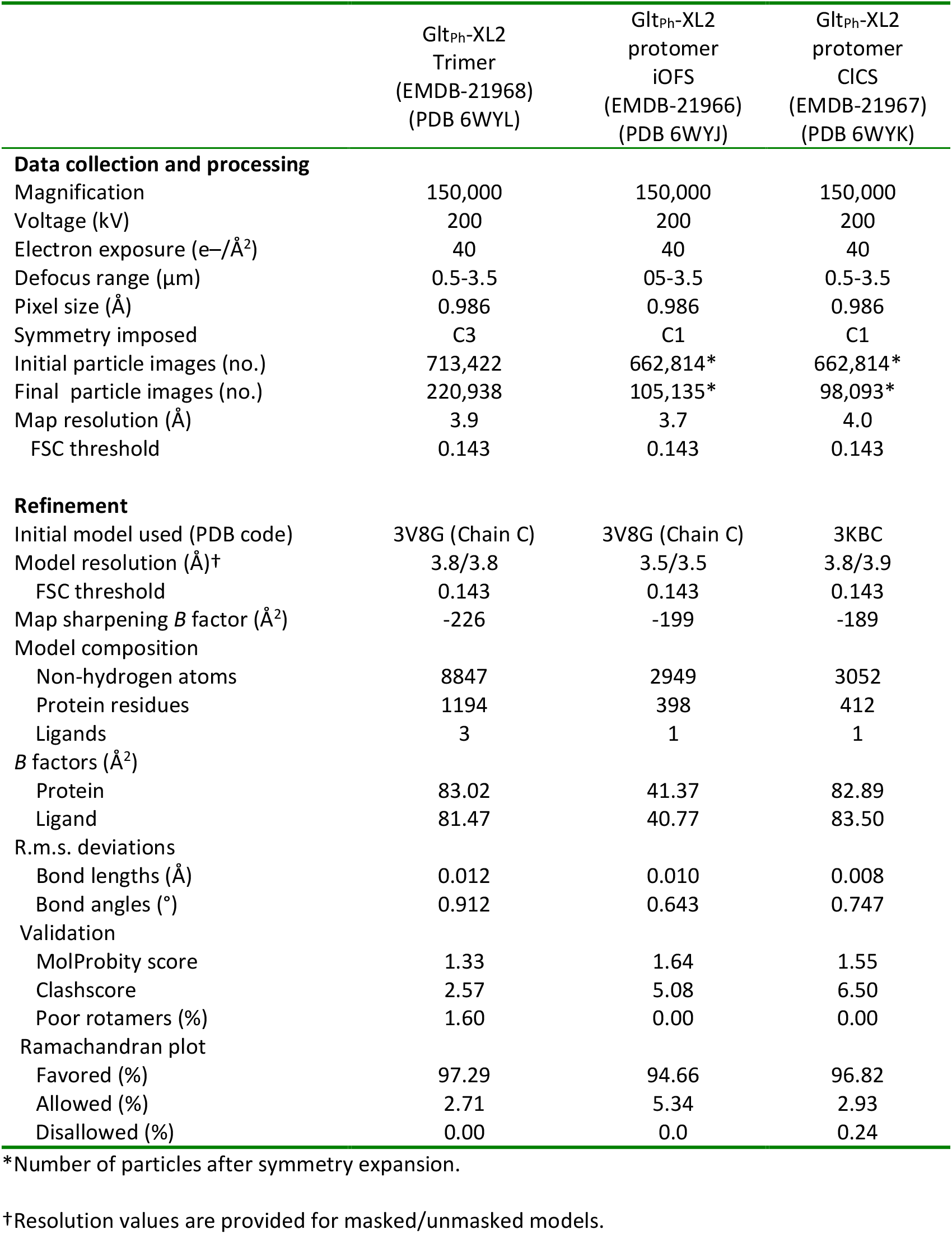
Cryo-EM data collection, refinement and validation statistics.

**Extended Data Table 3.** EAAT1 and corresponding GltPh residues that are within 5 Å of a permeating Cl^−^ ion.

| EAAT1 | Glt<sub>Ph</sub> | Location | Publication |
| --- | --- | --- | --- |
| R47 | E9 | TM1 | N/A |
| **F50<sup>c</sup>** | V12 | TM1 | 37 |
| V51 | L13 | TM1 | 37 |
| **T54<sup>c</sup>** | I16 | TM1 | 37 |
| Y75 | D37 | TM2 | N/A |
| K79 | T41 | TM2 | N/A |
| **L88<sup>c</sup>** | F50 | TM2 | 36, 37 |
| **M89<sup>c</sup>** | V51 | TM2 | 36 |
| R90 | R52 | TM2 | 34 |
| L92 | L54 | TM2 | 37 |
| Q93 | K55 | TM2 | 34 |
| **L99<sup>a</sup>** | I61 | TM2 | 34 |
| **S103<sup>a</sup>** | S65 | TM2 | 20, 34 |
| M279 | Y195 | TM5 | 20, 36 |
| R280 | K196 | TM5 | 36 |
| A283 | N199 | TM5 | N/A |
| **M286<sup>c</sup>** | M202 | TM5 | 36, 37 |
| W287 | Q203 | TM5 | N/A |
| **A289<sup>c</sup>** | A205 | TM5 | 37 |
| I293 | V209 | TM5 | 36, 37 |
| **L296<sup>b,c</sup>** | L212 | TM5 | 36, 37 |
| K300 | V216 | TM5 | N/A |
| E303 | E219 | TM5 | N/A |
| T358 | T271 | HP1 | 37 |
| G361 | V274 | HP1 | 37 |
| T362 | T275 | HP1 | 37 |
| S363 | R276 | HP1 | 29, 37 |
| S364 | S277 | HP1 | N/A |
| **S366<sup>a</sup>** | S279 | HP1 | 28 |
| A367 | G280 | HP1 | N/A |
| **L369<sup>a</sup>** | L282 | HP1 | 28 |
| P370 | P283 | HP1 | N/A |
| I371 | V284 | HP1 | 37 |
| F373 | M286 | HP1 | 20, 28, 36 |
| K374 | R287 | HP1 | N/A |
| **P392<sup>a</sup>** | P304 | TM7 | 28, 36 |
| A395 | A307 | TM7 | N/A |
| T396 | T308 | TM7 | 28 |
| P444 | P356 | HP2 | N/A |
| **Q445<sup>b</sup>** | G357 | HP2 | N/A |
| G447 | G359 | HP2 | N/A |
| L448 | A360 | HP2 | N/A |
| **R477<sup>b</sup>** | M391/R276* (HP1) | TM8 | 29, 36 |

Residues in the S103<sup>a</sup> and the R477<sup>b</sup> sites. Hydrophobic gate residues mutated in this study<sup>c</sup>
\*R477 of EAAT1 corresponds to R276, found in HP1 of Glt<sub>Ph</sub>

## Acknowledgements

This work was supported by the Australian National Health and Medical Research Council Project Grant APP1164494 (RR, RV, JF) and Fellowship APP1159347 (AGS), National Institutes of Health grants P41-GM104601 (ET) and R01-GM067887 (ET), computational resources provided by XSEDE (grant MCA06N060 to ET), Research Training Program Scholarship (IC) and Beckman Institute Graduate Fellowship (SP).

We wish to thank and acknowledge the use of the Victor Chang Cardiac Research Institute Innovation Centre, funded by the NSW Government, and the Electron Microscope Unit at UNSW Sydney, funded in part by the NSW Government; and the use of the MX2 beamline at the Australian Synchrotron, part of ANSTO, and the Australian Cancer Research Foundation (ACRF) detector. We acknowledge the facilities and technical assistance of Microscopy Australia at the Australian Centre for Microscopy & Microanalysis at the University of Sydney. We would also like to thank Xiaoyu Wang and Olga Boudker for helpful discussions, Zhiyu Zhao for his assistance with simulations, and Cheryl Handford and those that support the *Xenopus laevis* colony at the University of Sydney.

## Author contributions

IC, JF, RC, RR designed and performed biochemistry and X-ray crystallography experiments. IC, JF, MS, AS designed and performed cryo-EM experiments. SP and ET designed the simulation experiments, SP performed and analyzed the simulations. QW, RC, RV, RR designed and performed functional experiments in oocytes. The manuscript was written by IC, SP, QW, RR with input from all authors.

## Materials & Correspondence

Correspondence to Renae Ryan.

